# Human ANKLE1 is a nuclease specific for branched DNA

**DOI:** 10.1101/2020.07.03.186551

**Authors:** Junfang Song, Alasdair D. J. Freeman, Axel Knebel, Anton Gartner, David M. J. Lilley

## Abstract

All physical connections between sister chromatids must be broken before cells can divide, and eukaryotic cells have evolved multiple ways in which to process branchpoints connecting DNA molecules separated both spatially and temporally. A single DNA link between chromatids has the potential to disrupt cell cycle progression and genome integrity, so it is highly likely that cells require a nuclease that can process remaining unresolved and hemi-resolved DNA junctions and other branched species at the very late stages of mitosis. We argue that ANKLE1 probably serves this function in human cells (LEM-3 in *C. elegans*). LEM-3 has previously been shown to be located at the cell mid-body, and we show here that human ANKLE1 is a nuclease that cleaves a range of branched DNA species. It thus has the substrate selectivity consistent with an enzyme required to process a variety of unresolved and hemi-resolved branchpoints in DNA. Our results imply that ANKLE1 acts as a catch-all enzyme of last resort that allows faithful chromosome segregation and cell division to occur.

## INTRODUCTION

To ensure faithful genome maintenance, chromatids must be precisely segregated to daughter cells. This requires the removal of all physical connections between sister chromatids before cells divide. These include intermediates of DNA recombination, such as four-way (Holliday) junctions, points at which chromatids become intertwined, and loci that have not been replicated by the time cells reach the metaphase-anaphase transition. Cytologically, DNA connections between segregating chromatids take the form of chromatin bridges (1-3), or ultrafine bridges (4,5); the latter do not stain with DAP1 but are associated with BLM and PICH helicases. DNA bridges form in each cell cycle, their number being increased under DNA damaging conditions, or when DNA replication is challenged (6). Failure to process DNA bridges impedes the segregation of chromatids and leads to genome instability via chromosome breakage, or cleavage furrow regression during cytokinesis resulting in binucleated cells and polyploidy.

To ensure that all connections are removed before cells divide, cells have evolved highly redundant mechanisms that remove all bridges. In eukaryotes four-way helical (Holliday) junctions are processed during S-phase by dissolution, using the combined activities of the Blooms helicase and topoisomerase IIIα (7-9). If junctions persist they are resolved by two major junction-cleavage activities using structure-specific nucleases. The SLX4-SLX1-MUS81-EME1 hetero-tetramer acts in the nucleus and is part of a larger complex of nucleases organised by the SLX4 scaffold protein (10-13). Within this complex SLX1 and MUS81 are structure-specific nucleases. Remaining junctions are processed by the cytoplasmic GEN1 (Yen1 in budding yeast) nuclease that acts during late M-phase and anaphase (14-16). GEN1 function as a homodimer, resolving junctions by two sequential cleavage reactions (17-19). The properties of GEN1 are very similar to junction-resolving enzymes of lower organisms and phage (20).

There is considerable redundancy between junction dissolution and resolution activities, and genetic interaction between the corresponding mutants have been reported. The loss of a single activity is generally well tolerated, but when cells become defective in multiple activities then they become very sensitive to DNA damage (21-23) and aberrant chromosomes are formed (23-25). It has been shown in human cells that the combinations of SLX4 and either BLM or GEN1 are synthetic lethal (26), underlining the vital importance of having a functional junction resolution activity.

Despite this considerable redundancy, a single DNA link between chromatids has the potential to disrupt cell cycle progression and genome integrity, and it is likely that cells require a nuclease that can process remaining unresolved and hemi-resolved DNA junctions and other branched species at the very late stages of mitosis, and a contender for this activity is LEM-3/ANKLE1. LEM-3 was discovered in a genetic screen for embryonic lethality in *Caenorhabditis elegans* following ionizing irradiation (27). The protein sequence has a predicted GIY-YIG nuclease domain, and was shown to exhibit nucleolytic activity on supercoiled DNA. ANKLE1 is the corresponding ortholog in humans and other metazoans (Figure S1) (28). Several lines of evidence now suggest that LEM-3/ANKLE1 might be responsible for processing DNA bridges during anaphase. First, LEM-3 has been found to interact genetically with MUS81, SLX1 and SLX 4 with embryonic lethality resulting in *lem*-3 double mutants, all single mutants being viable (3,29). Second, LEM-3 is excluded from the nucleus and accumulates at the mid-body the structure where the two daughter cells finally abscise from each other, starting from the early stages of cytokinesis, and genetic evidence indicate that such localisation is important for LEM-3 activity *in vivo* (30). Finally, chromosomal bridges persist in *lem*-3 mutants upon treatment with various DNA damaging agents, and under conditions where DNA replication or chromosome decondensation is compromised (30). Thus LEM-3/ANKLE1 exhibits many of the properties that would be expected for a nuclease whose role is to process persistent helical junctions that would otherwise lead to the regression of the cytokinetic furrow and ensuing polyploidisation and aneuploidy.

What is currently missing from this analysis is a knowledge of the substrate specificity for either LEM-3 or ANKLE1. We have therefore expressed human ANKLE1 in insect cells and studied its nuclease activity on a wide variety of branched DNA species. We find that ANKLE1 is a nuclease that is specific for branched DNA, cleaving a wide range of helical junctions, flap structures and splayed Y-junctions. Thus ANKLE1 has exactly the type of substrate specificity that would be expected for an enzyme that is located at the cell mid-body whose role is to act as a catch-all final defence to process DNA junctions that have evaded all the other levels of enzymatic attack. Our results imply that ANKLE1 (and most probably LEM-3) acts as the enzyme of last resort allowing for faithful chromosome segregation and cell division.

## RESULTS

### Expression of human ANKLE1 in insect cells

A synthetic cDNA encoding human ANKLE1 was constructed with codons optimized for expression in insect cells, carrying an N-terminal fusion with glutathione S-transferase (GST) and an intervening TEV protease cleavage site (Figure 1). The gene was incorporated into the baculovirus genome in *E. coli* DH10EMBacY by Tn7 transposition (31), and the resulting DNA was transfected into Sf9 insect cells. In order to obtain generation 2 recombinant virus (V_2_), two subsequent infections were carried out. Optimal production of hANKLE1 was achieved by infecting 1.0 x10^6^ cells/ml of freshly diluted Sf9 cells with 1:500 (v/v) V_2_. GST-tagged hANKLE1 was purified using affinity chromatography on glutathione Sepharose 4B, followed by TEV cleavage on the column to remove the GST tag. The resulting non-fusion hANKLE1 was further purified by gel filtration after which the hANKLE1 migrated as a single species at the expected size on polyacrylamide gel electrophoresis in the presence of SDS (Figure 1). The protein was confirmed as human ANKLE1 by fragmentation-mass spectrometric analysis of trypsin-digested protein (Figure S2). Two isoforms of hANKLE1 were investigated with or without an N-terminal extension of 54 amino acids (Figure S1). Both forms (with and without an N-terminal GST tag) were found to have similar nuclease activity on branched DNA (Figure S3). The majority of experiments here were performed using the 615 amino acid form with the GST tag removed.

**Figure 1.**
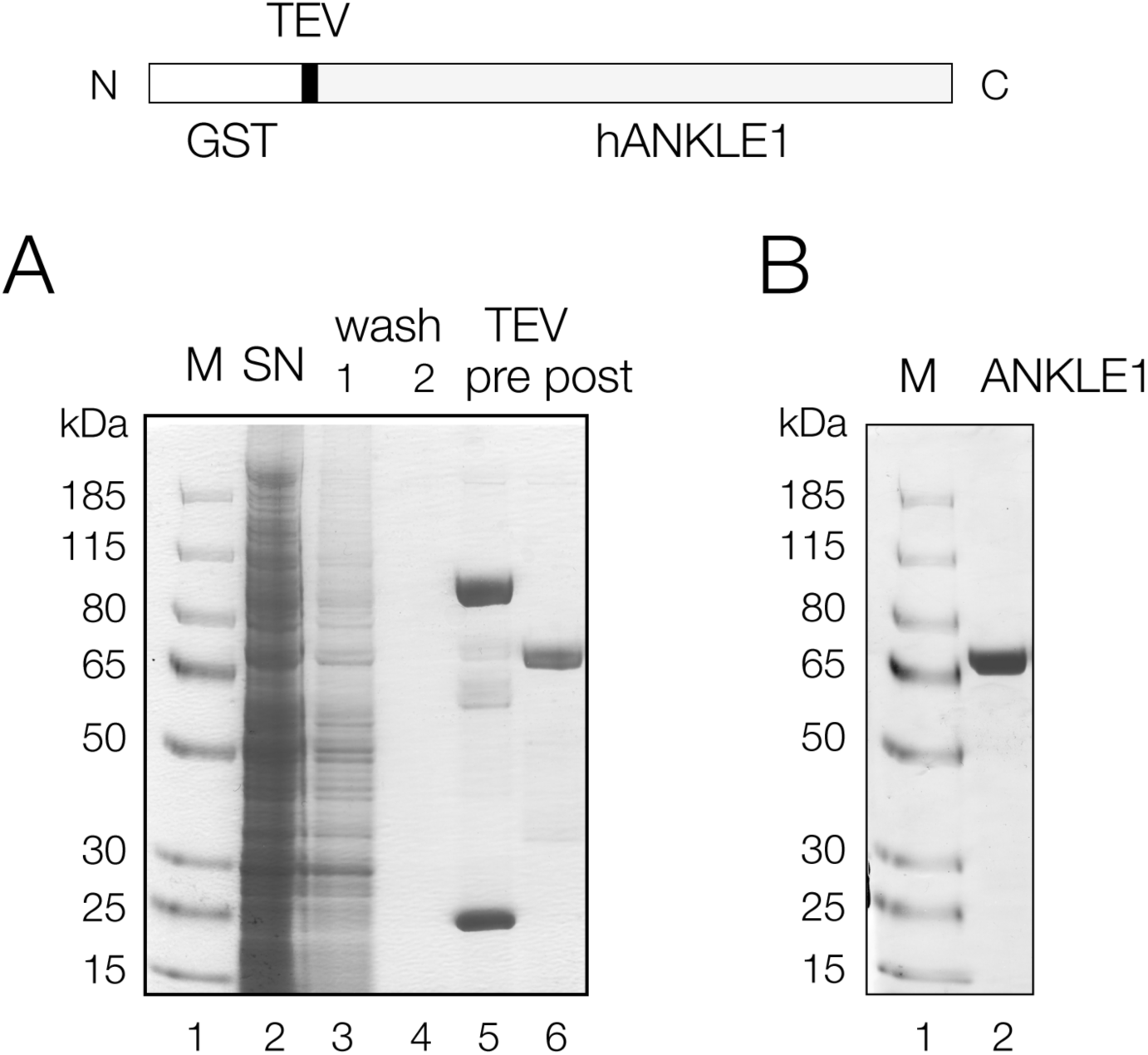
Expression and purification of human ANKLE1 in insect cells. Top. Schematic of GST-hANKLE1 fusion protein. The fusion can be cleavage by TEV protease to release hANKLE1. **A**. Denaturing gel electrophoresis of expressed hANKLE1. Sf9 insect cells were lysed and centrifuged. The soluble fraction was incubated with glutathione Sepharose 4B for one hour, allowed to settle and the supernatant taken (track 2). The resin was treated with TEV protease. Track 1. protein markers, molecular masses are indicated left ; track 2. proteins not binding to the glutathione resin in cytoplasmic extract; tracks 3, 4, sequential washes of the glutathione resin ; track 5. proteins binding to the glutathione resin in cytoplasmic extract, including GST-hANKLE1, endogenous insect GST, and some proteins whose sizes are smaller than GST-hANKLE1 (these could include incomplete translation products) ; track 6. hANKLE1 released from the resin after cleavage with TEV protease. **B**. Denaturing gel electrophoresis of unfused hANKLE1 after gel filtration. Track 1. protein markers, molecular masses are indicated left ; track 2. purified hANKLE1.

### Human ANKLE1 is a nuclease with specificity for branched DNA

We first explored whether or not human ANKLE1 expressed and purified from insect cells exhibits nuclease activity on branched DNA species. The sequences of the DNA substrates were all derived from a well-characterized four-way DNA junction 3 (32) (the provenance of all the junctions is explained in Figure S4), each with a radioactive-[5’-^32^P] label on the same x strand. We found that a variety of three- and four-way DNA junctions, flap and needle structures were cleaved by hANKLE1 (Figure 2, Figure S5). We observed that the enzyme was especially active in the presence of Mn^2+^ ions (Figure S6), and optimal conditions for cleavage were found to be 20 mM cacodylate (pH 6.5), 50 mM KCl, 2 mM MnCl_2_, 0.1mg/ml BSA. 5 nM DNA substrate were incubated with 330 nM hANKLE1 i.e. experiments were performed under single-turnover conditions.

**Figure 2.**
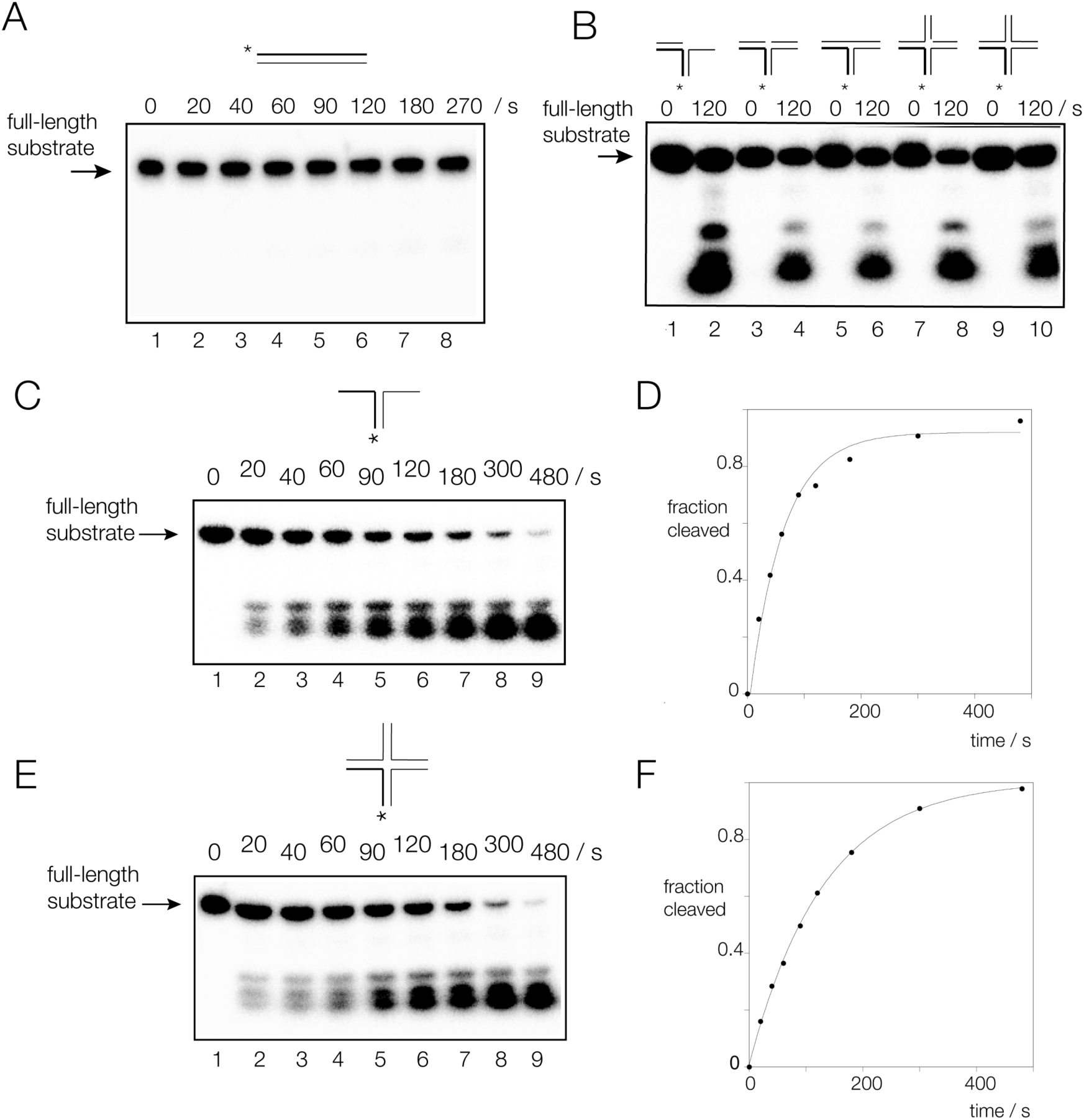
Human ANKLE1 selectively cleaves a variety of branched DNA species. A series of DNA species were formed by hybridization of component strands, all containing the same radioactively [5’- ^32^P]-labeled x strand indicated by an asterisk. The provenance of these species is shown in Figure S4 and their sequences in Table S1. DNA was incubated with hANKLE1 under single turnover conditions in 20 mM cacodylate (pH 6.5), 2 mM MnCl_2_, 50 mM KCl, 0.1 mg/ml BSA at 37°C. After the indicated times of incubation the reaction was terminated and the substrate and products separated by denaturing gel electrophoresis. **A**. Duplex DNA. The DNA was incubated with hANKLE1 for 270 s, with aliquots removed at the indicated times. **B**. Five different branched DNA species were incubated with hANKLE1 for 0 and 120 s, and cleavage analysed by denaturing gel electrophoresis. The species were: tracks 1 and 2, a 5’ flap structure; tracks 3 and 4, a three-way junction with a nick at the point of stand exchange; tracks 5 and 6, an intact three-way junction; tracks 7 and 8, a four-way junction with a nick at the point of stand exchange; an intact four-way junction. **C, D**. A splayed Y_X_-junction. The DNA was incubated with hANKLE1 for 480 s, with aliquots removed at the indicated times. Note that the substrate is almost fully cleaved by the end of the incubation. Part **D** shows a plot of reaction progress as a function of time. The data (points) were fitted to a single exponential function (line). **E, F**. A four-way (Holliday) junction. The DNA was incubated with hANKLE1 for 480 s, with aliquots removed at the indicated times. The junction is also almost fully cleaved by the end of the incubation. Part **F** shows a plot of reaction progress as a function of time. The data (points) were fitted to a single exponential function (line).

hANKLE1 exhibits low activity on duplex DNA (Figure 2A) or single-stranded DNA (data not shown), but is active on a wide variety of branched DNA (Figure 2B-D, Figure S4). Time courses of incubation of a splayed Y_X_ junction (i.e. a duplex with non-complementary single-stranded extensions) and a 4H four-way junction show that both are cleaved to completion. Fitting reaction progress to single exponential functions gives rates of *k*_obs_ = 0.016 and 0.008 s^-1^ for the Y-junction and four-way junctions respectively (Figure 2C, D), and the splayed Y_X_ junction is cleaved slightly more efficiently than the other substrates. We observe that a splayed Y_H_ junction (this is derived from the top half of junction 3, see Figure S4) is cleaved to completion (Figure S7), yet is totally unrelated to the splayed Y_X_ junction in terms of sequence. We conclude that human ANKLE1 is primarily a structure-selective nuclease of broad specificity capable of cleaving a wide variety of branched DNA species.

### ANKLE1 cleaves double-stranded DNA close to branchpoints

We next mapped ANKLE1 cleavage sites in the different branched DNA substrates at the nucleotide level (Figure 3A). These are four-way (4H) and three-way (3H) helical junctions, a three-way junction with a single-strand break at the point of strand exchange, 5’ and 3’ flaps and a splayed Y_X_ junction comprising a duplex with non-complementary single stranded sections at one end. Each species can be derived from the four-way junction 3 (see Figure S4), and include a common strand (the x strand) that was radioactively [5’-^32^P]-labelled. This strand is cleaved at the same sites in each branched DNA species by ANKLE1. All the cleavage occurs 5’ to the point of strand exchange; in double-stranded DNA in each case, including the splayed Y-junction. Major cleavages are observed 3, 9 and 14 nucleotides 5’ to the point of strand exchange; thus these sites have been designated by these numbers. Each site conforms to the sequence TRi (R = purine), indicating a degree of sequence selectivity in cleavage. However the primary selectivity is at the level of structure – these sites must be adjacent to a junction for optimal cleavage.

**Figure 3.**
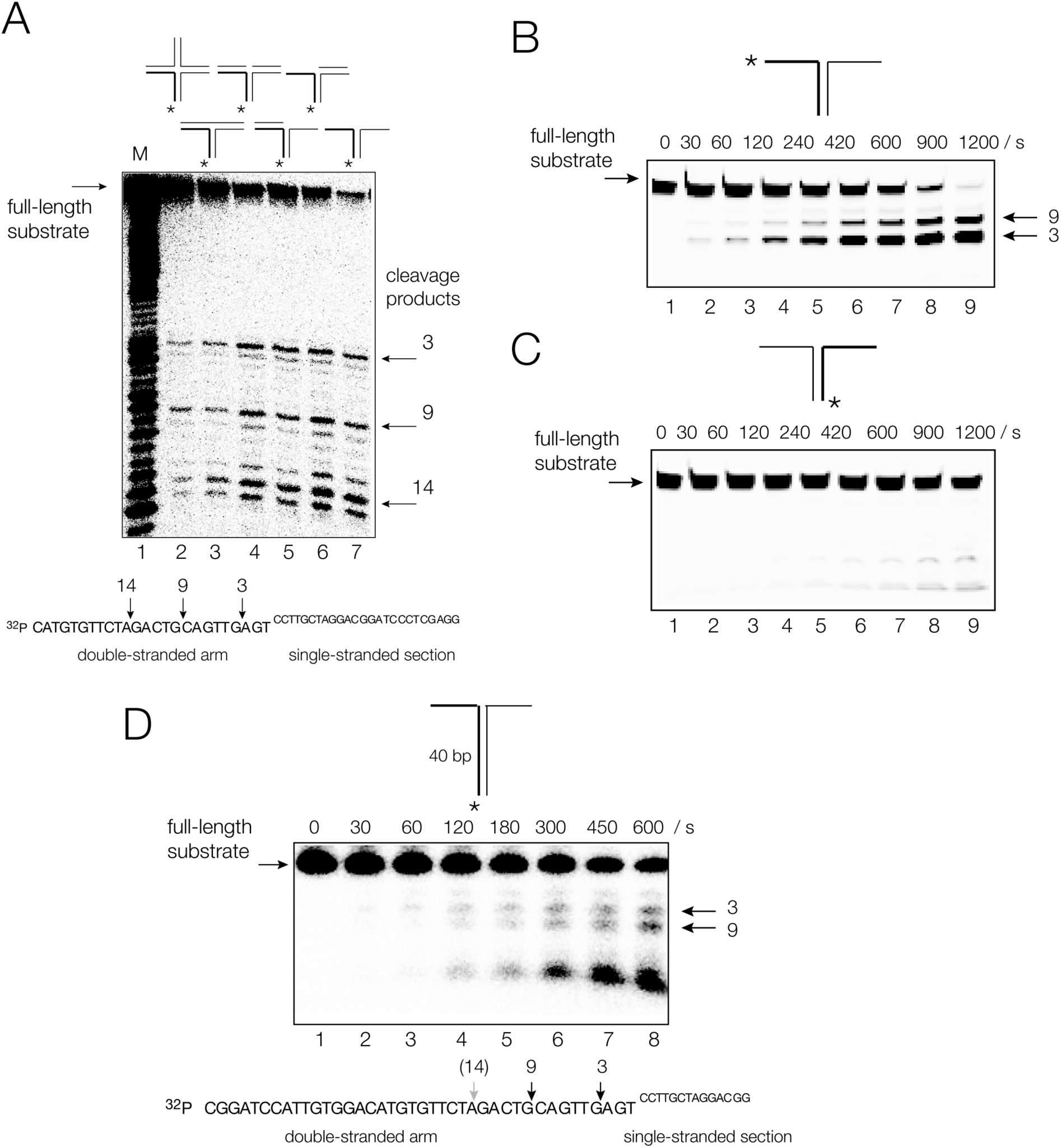
Human ANKLE cleaves double-stranded DNA close to helical branchpoints. **A**. Cleavage positions in a series of branched DNA species analysed by denaturing gel electrophoresis at single-nucleotide resolution. The positions of cleavage are arrowed on the sequence of the radioactively [5’-^32^P]-labeled x strand shown below the autoradiograph. **B**. Analysis of the cleavage of the splayed Y_X_-junction with the x strand fluorescently labelled at its 3’ end. The DNA was incubated with hANKLE1 for 1200 s, with aliquots removed at the indicated times. This reveals that the major cleavage occurs at site 3, i.e. 3 nt from the point of strand exchange. **C**. Analysis of the cleavage of the splayed Y_X_-junction with the r strand fluorescently labelled at its 3’ end. The DNA was incubated with hANKLE1 for 1200 s, with aliquots removed at the indicated times. Note that the r strand is very weakly cleaved. **D**. Cleavage of a splayed Y_X_-junction with an extended double-stranded helical arm of 40 bp. Analysis of the cleavage of the splayed Y-junction with the x strand radioactively [5’-^32^P]-labeled. The DNA was incubated with hANKLE1 for 600 s, with aliquots removed at the indicated times.

The cleavages on the splayed Y_X_ junction were investigated in greater detail. First, the same x strand was fluorescently-labelled at its 3’ end by fluorescein. ANKLE1 cleaves the strand to completion, cleaving site 3 twice as fast as at site 9 (Figure 3B). In this experiment cleavage at site 14 was not observed, indicating that it occurs subsequently to cleavage at sites 3 or 9. Second, we fluorescently 3’-labelled the other (r) strand of the junction. By comparison with the x strand there is very little cleavage on the r strand by ANKLE1 (Figure 3C). Finally we constructed a version of the splayed Y_X_ junction in which the double-stranded section was extended to 40 bp in length. Incubation with ANKLE1 lead to cleavage at a new site closer to the 5’ end of the double-stranded arm. We conclude that enzyme predominantly cleaves on DNA strand 5’ to a helical junction. The significant cleavage sites are the 3 and 9 sites, but additional cleavages occur close to the 5’ end of the molecule. These more distant sites are probably a result of having an open end and are probably not relevant to the normal function of ANKLE1.

### The GIG-YIG motif of human ANKLE1 is required for nuclease cleavage close to DNA branchpoints

ANKLE1 has a highly conserved GIY-YIG domain at its C-terminal end (Figure 4A). GIY-YIG domains function as the active centers of a number of nucleases (33), including R-*Eco*29kl (34), I-TevI (35), UvrC (36) and the junction-resolving enzyme SLX1 (37). For ANKLE1 the sequence is FTY_453_ (31 amino acids) Y_486_VG. The tyrosine residues play an important role in the probable mechanism of phosphodiester bond hydrolysis, so we made a Y453F mutation that removes the phenolic hydroxyl of the tyrosine. The resulting atomic mutant of ANKLE1 was essentially inactive on the splayed Y-junction (Figure 4B), showing the importance of the GIY-YIG domain in the function of the nuclease. In confirmation of this we have also found that a Y_486_F mutant was similarly inactive (Figure S8). We conclude that both tyrosine phenolic oxygen atoms of the GIY_YIG motif are essential for the cleavage of DNA junctions by human ANKLE1.

**Figure 4.**
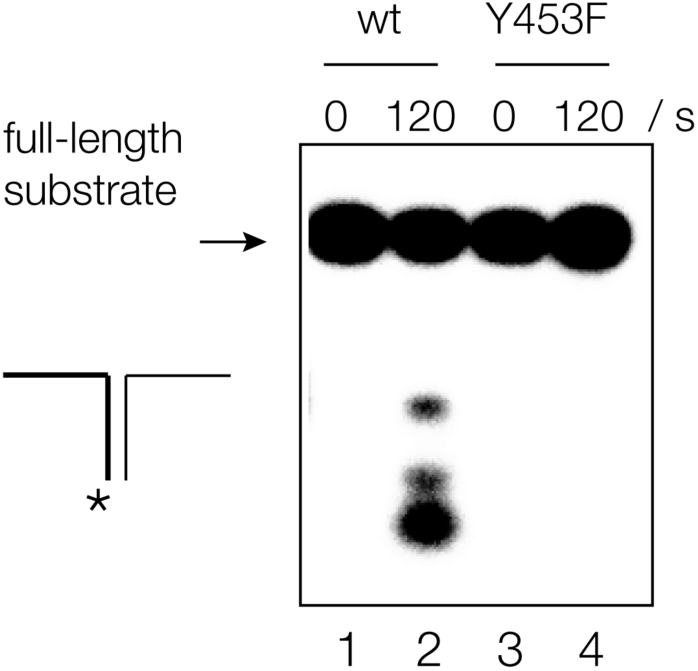
Cleavage of branched DNA by human ANKLE requires the GIY-YIG domain. A mutant in the first tyrosine residue of the GIY-YIG domain (Y453F) was prepared. A splayed Y_X_-junction radioactively [5’-^32^P]-labeled on the x strand was incubated with either wild-type hANKLE1 (tracks 1 and 2) or Y453F hANKLE1 (tracks 3 and 4) for either 0 or 120 s. Note that the DNA incubated with the mutant enzyme was uncleaved. Note that only a single atom of hANKLE1 has been changed by the mutation. A corresponding mutant in the other tyrosine (Y486F) was also inactive (Figure S8).

## DISCUSSION

These studies have shown that human ANKLE1 expressed in insect cells is a nuclease that is selective for branched DNA species. Helical junctions with and without single-strand breaks, flaps of various kinds and splayed Y-structures are all subject to hydrolysis of phosphodiester linkages at sites in double-stranded DNA close to the junction. The range of branched DNA substrates is wider than a conventional Holliday junction-resolving enzyme such as GEN1, consistent with a role in processing unresolved or partially resolved junctions that might result from a failure to complete processing of recombination intermediates. Resolving enzymes like GEN1 have evolved processes whereby the second strand cleavage event is significantly accelerated relative to the first (17,38,39), so increasing the probability of bilateral cleavage during the lifetime of the complex. However, if that fails for any reason then a hemi-resolved junction could result. A number of different branched species might result from aberrant processing of junctions, any of which might result in DNA bridging between chromatids. It is therefore probable that a nuclease with relatively wide substrate specificity would be required to clean up debris from these events.

Fluorescence microscopy has previously shown that the *C. elegans* LEM-3 that is orthologous with ANKLE1 (Figure S1) is located at the cell mid-body, where it likely contributes to the resolution of residual DNA bridges just before cells divide (30). These bridges can be induced by ionizing radiation and mutagens such as hydroxyurea, by slowing DNA replication, by DNA de-condensation defects, as well as by inhibition of enzymes associated with Holliday junction dissolution or resolution such as MUS81, SLX1 and SLX4 (3). While the exact nature and origins of DNA bridges are not fully understood at present, they are observed from the onset of anaphase, through telophase, up to the point where cells divide during cytokinesis and it is likely that they arise in part from failed or incomplete resolution of recombination intermediates, that must be processed before cell division can be completed. We have now shown that human ANKLE1 has broad-range structural selectivity for a range of branched DNA species. That together with the sub-cellular location of the closely-related LEM3 in *C. elegans* is fully consistent with a role at cytokinesis to process DNA branchpoints that have not been resolved at an earlier stage of the cell cycle. The broad specificity could lead to the cleavage of branched DNA structures that occur during DNA duplication, and overexpression of ANKLE1 in the nucleus was shown to trigger DNA damage (28). This might explain why LEM-3 appears to be excluded from the nucleus (3). Put briefly ANKLE1 has the required specificity and the correct location to act as a catch-all “enzyme of last resort” to process a variety of junctions in residual DNA bridges in order to allow cell division to be completed.

## MATERIALS AND METHODS

### Expression and purification of human ANKLE1

DNA with the human ANKLE1 coding sequence, codon optimized for baculovirus expression was synthesized by GeneArt Gene Synthesis (Thermofisher). The DNA was cloned into MultiBac expression vector pKL with an N-terminal GST tag. Active site mutants Y453F and Y486F were generated by site-directed mutagenesis (QuikChange, Agilent). DNA encoding wild type or mutant hANKLE1 was integrated into baculovirus genome by the transformation into *E*.*coli* DH10EMBacY by Tn7 transposition. The recombinant bacmids, initial DNA (V_0_), generation 1 (V_1_) and 2 virus (V_2_) containing ANKLE1 were prepared according to protocols published by Berger et.al. (31).

Insect Sf9 cells at a density of 10^6^ cells/ml were infected with V_2_ virus in a 1:500 ratio (vol/vol) and incubated at 27°C shaking at 150 rpm. The cells were harvested at day 3 of proliferation arrest, the cell pellet was suspended and lysed in lysis buffer containing 20 mM HEPES (pH 7.5), 500 mM KCl, 10% glycerol, 1 mM DTT, 0.1% Triton-X and EDTA-free Protease Inhibitor Cocktail (cOmplete™, Roche). After centrifugation at 20,000 g at 4°C the cytoplasmic extract was incubated with glutathione Sepharose® 4B (GE) for 1 h at 4°C. The resin was washed with 20 mM HEPES (pH 7.5), 500 mM KCl, 10% glycerol, 1 mM DTT, 0.1% Triton-X. After washing, the resin was incubated with TEV Protease (NEB) for 3 h at room temperature. The cleaved hANKLE1 was then subject to gel filtration chromatography on Superdex^®^ 200 (GE). hANKLE1 concentration was estimated spectrophotometrically by absorption at 280 nm using an extinction coefficient of 66,000 M^-1^ cm^-1^.

### Characterisation of expressed human ANKLE1 by fragmentation and mass spectrometry

A fragment of polyacrylamide gel containing the human ANKLE1 was destained, and reacted with 10 mM DTT and 55 mM iodoacetamide to reduce disulfide bonds and alkylate free cysteines. The protein was then digested with 12.5 *µ*g/ml trypsin in 20 mM ammonium bicarbonate at 30°C for 16 h. The peptide mixture was dried and suspended to 50 *µ*l 1% formic acid, and then separated by reversed-phase chromatography. Peptides were initially trapped on an Acclaim PepMap 100 (C18, 100 *µ*M x 2 cm) and then separated on an Easy-Spray PepMap RSLC C18 column (75 *µ*M x 50 cm) (Thermo Scientific). Samples were transferred to a Q Exactive plus mass spectrometer via an Easy-Spray source with temperature set at 50°C and a source voltage of 2.0 kV. The precursor ion is selected by the quadrupole and isolated for fragmentation using higher energy collisional dissociation. The ions transferred to an orbitrap to provide high resolution accurate mass data of the fragmented peptides. The peptides are identified by the data analysis software by reference to a protein database.

### Oligonucleotide synthesis

Oligonucleotides were synthesised using β-cyanoethyl phosphoramidite chemistry {Beaucage, 1981 #6; Sinha, 1984 #171}. Fully deprotected oligonucleotides were purified by gel electrophoresis on a 10%(w/v) polyacrylamide gel in 90 mM Tris.borate (pH 8.5), 2 mM EDTA (TBE buffer) containing 8M urea. Oligonucleotides were detected by UV shadowing and recovered by electroelution and ethanol precipitation. Concentration estimated by absorbance at 280 nm. Oligonucleotides labelled with fluorescein at their 3’-termini were synthesized using 6-fluorescein CPG columns (Glen Research 20-2961). Oligonucleotides were radioactively [5’-^32^P]-labelled using [γ-^32^P] ATP (Perkin Elmer) using T4 polynucleotide kinase (Fermentas) for 30 min at 37°C in in 50 mM Tris (pH 7.6), 10 mM MgCl_2_, 5 mM DTT, 0.1 mM spermidine. All DNA sequences used are tabulated in Table S1.

### Preparation of DNA substrates

Substrates were hybridized by mixing 1 *µ*M of one radioactively [5’-^32^P]-labelled strand with 1.5 *µ*M of the other required strands (see Figure S4) and incubated for 2 min at 80°C followed by slow cooling overnight. DNA substrates were purified by electrophoresis in 6% (29:1) polyacrylamide gel in TBE. The labelled DNA was recovered by electroelution in 0.5 x TBE. After ethanol precipitation, the DNA substrate concentration was measured spectrophotometrically by absorption at 260 nm and adjusted to 50 nM. Splayed Y junctions with a 3’-fluorescently-labelled strand were prepared by mixing equimolar quantities of labelled and unlabelled strands at 20 *µ*M and hybridising and purified as above.

### Analysis of DNA cleavage by human ANKLE1

5 nM [5’-^32^P]-labelled DNA substrate was incubated with 330 nM hANKLE1 for 10 min at 20°C in 20 mM cacodylate (pH 6.5), 50 mM KCl, 0.1 mg/ml BSA. After a 3 min pre-incubation at 37°C, the cleavage reaction was initiated by the addition of MnCl_2_ (unless stated otherwise) to a final concentration of 2 mM. Aliquots were taken at chosen times and the reaction terminated by addition of EDTA to a final concentration of 20 mM and 50% formamide. The aliquots were heated at 95°C for 15 min and then loaded on to a 14% (19:1) polyacrylamide gel in TBE, 8M urea. Radioactive DNA was detected by exposure to a storage phosphor screens, and visualised using a BAS-1500 phosphoimager (Fuji). Fluorescent DNA was visualised in the gel using a Typhoon FLA9000 imager using a 473 nm laser. Data were analysed as the fraction of cleaved DNA (*f*_c_) versus time (*t*) and fitted by non-linear regression to :

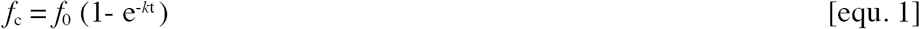

where *f*_0_ is the fraction of DNA cleaved at the end of the reaction and *k* is the observed rate of cleavage.

## ACKNOWLEGEMENTS

We thank Drs Anne-Cécile Déclais and Tim Wilson for discussion, and Kenny Beattie, Samantha Kosto and Douglas Lamont for mass spectrometry data collection and analysis. We are grateful to Elisa Garcia-Wilson, Taciana Kasciukovic and Harinath Doodhi for technical assistance in baculovirus expression system and protein purification. We acknowledge Cancer Research UK (program grant A18604) and BBSCRC (BB/S002782/1) for financial support of our research in Dundee, and the Korean Institute for Basic Science (IBS-R022-A1-2019).

