## Supplemental Information for "Human ANKLE1 is a nuclease specific for branched DNA"

### SUPPLEMENTARY INFORMATION

#### SUPPLEMENTARY FIGURES

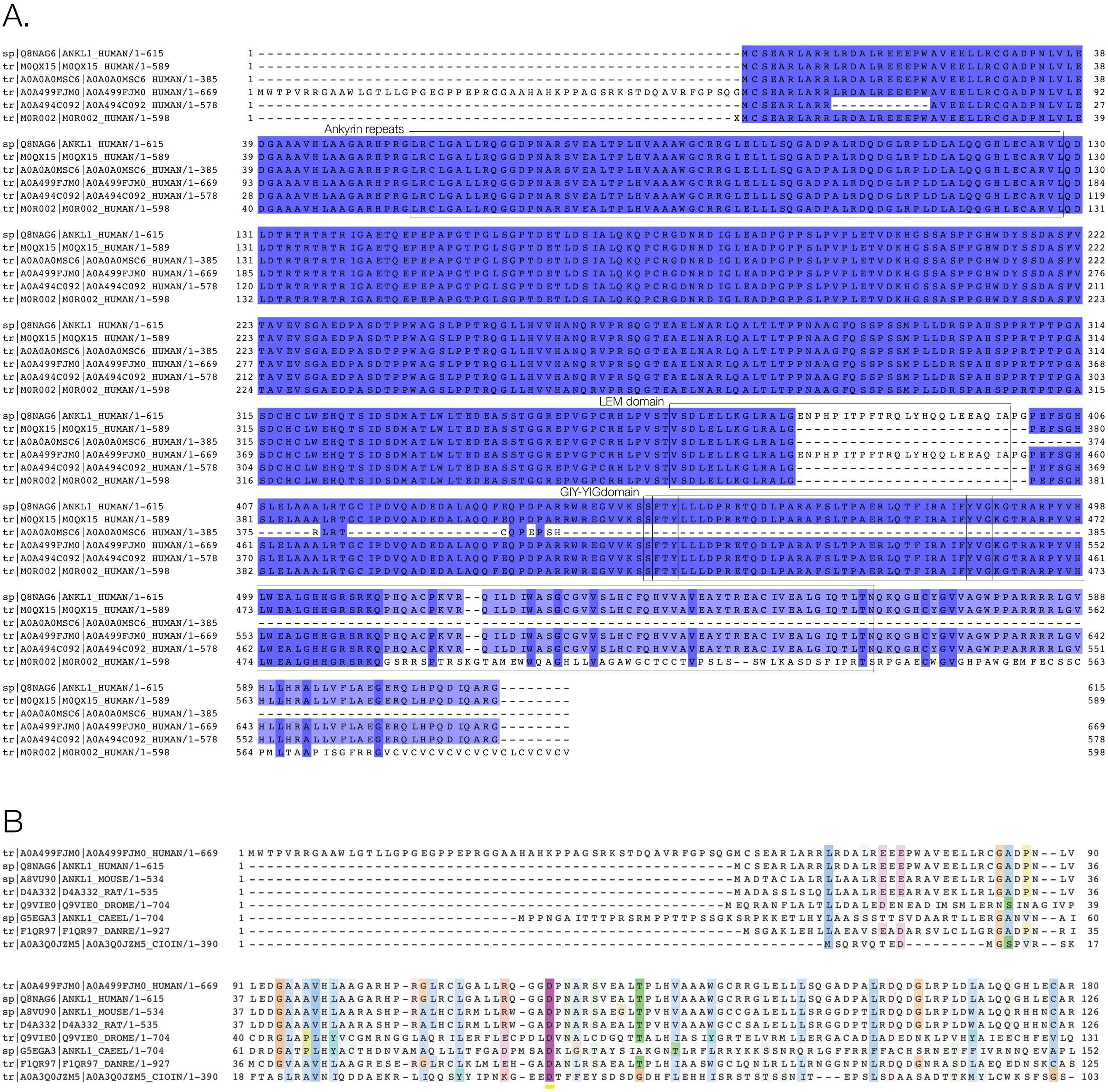

**Appendix Figure S1.** Alignment of human isoforms of ANKLE1.

**A**. The sequences of six human ANKLE1 isoforms were obtained from the UniProt database. Multiple sequence alignment was performed using Clustal via Jalview. The alignment shows that two isoforms Q8NAG6 and A0A499FJM0 contain three functional domains highly conserved in ANKLE1 homologs in vertebrate, N-terminal Ankyrin repeats, LEM domain and a GIY-YIG domain (all boxed). A0A499FJM0 comprises 669 amino acids, and shares 100% sequence identity with Q8NAG6 (615 aa), except for an extension of 54 amino acids in its N-terminus.

**B**. Alignment of the N-terminal of ANKLE1 sequences from other species suggests the extra 54 amino acid residues in A0A499FJM0 are not present in all other species, from *C. elegans* to mouse. It therefore indicates that extra N-terminus sequence in Q8NAG6 may be not required activity. The majority of experiments here were performed using Q8NAG6, the 615 amino acid form.

```
  1  MCSEARLARR  LRDALREEEP  WAVEELLRCG  ADPNLVLEDG  AAAVHLAAGA
 51  RHPRGLRCLG  ALLRQGGDPN  ARSVEALTPL  HVAAAWGCRR  GLELLLSQGA
101  DPALRDQDGL  RPLDLALQQG  HLECARVLQD  LDTRTRTRTR  IGAETQEPEP
151  APGTPGLSGP  TDETLDSIAL  QKQPCRGDNR  DIGLEADPGP  PSLPVPLETV
201  DKHGSSASPP  GHWDYSSDAS  FVTAVEVSGA  EDPASDTPPW  AGSLPPTRQG
251  LLHVVHANQR  VPRSQGTEAE  LNARLQALTL  TPPNAAGFQS  SPSSMPLLDR
301  SPAHSPPRTP  TPGASDCHCL  WEHQTSIDSD  MATLWLTEDE  ASSTGGREPV
351  GPCRHLPVST  VSDLELLKGL  RALGENPHPI  TPFTRQLYHQ  QLEEAQIAPG
401  PEFSGHSLEL  AAALRTGCIP  DVQADEDALA  QQFEQPDPAR  RWREGVVKSS
451  FTYLLLDPRE  TQDLPARAFS  LTPAERLQTF  IRAIFYVGKG  TRARPYVHLW
501  EALGHHGRSR  KQPHQACPKV  RQILDIWASG  CGVVSLHCFQ  HVVAVEAYTR
551  EACIVEALGI  QTLTNQKQGH  CYGVVAGWPP  ARRRRLGVHL  LHRALLVFLA
601  EGERQLHPQD  IQARG
```

**Appendix Figure S2.** Mass spectrometric characterisation of human ANKLE1 expressed in insect cells. The band from the polyacrylamide gel shown in Figure 1B was excised and analysed by peptide fragmentation and mass spectrometry. Peptides matching the sequence of hANKLE1 are colored red; these correspond to 72% of the total hANKLE1 protein sequence.

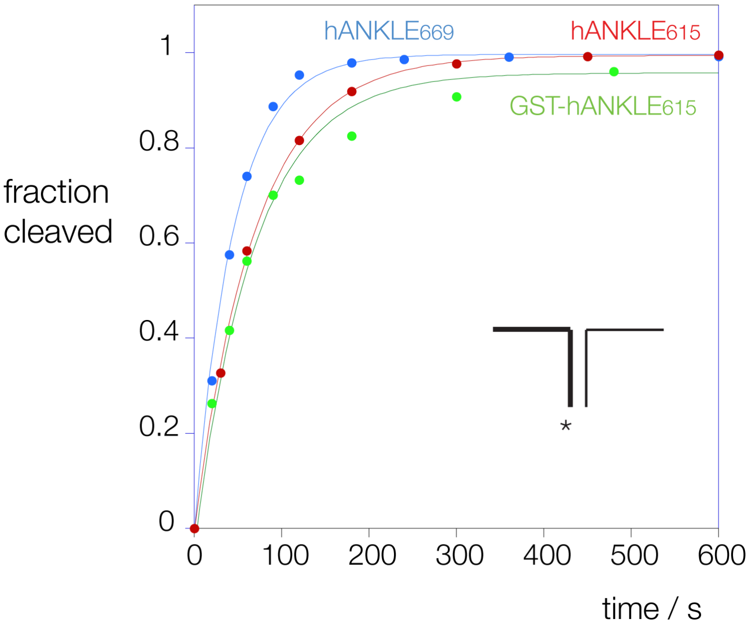


**Appendix Figure S3.** Reaction progress plotted for cleavage of splayed $Y_X$ junction by three forms of human ANKLE1. The three forms are the 669 amino acid polypeptide (blue), and the 615 amino acid polypeptide with (green) and without (red) an N-terminal GST peptide. Each reaction proceeds with a similar rate and each achieved near-full conversion to cleaved product. Fitting to single exponential functions (lines) gives rates of unfused 615 aa $k_{obs} = 0.015$ s$^{-1}$, GST-615 aa $k_{obs} = 0.015$ s$^{-1}$, and 669 aa $k_{obs} = 0.022$ s$^{-1}$. Reactions were performed under single-turnover conditions in 20 mM cacodylate (pH 6.5), 50 mM KCl, 2 mM $MnCl_2$, 0.1mg/ml BSA. The insert represents the structure of the $Y_X$ junction, in the same manner as the depictions in Figure S4. In this and other figures the asterisk shows the labelled terminus.

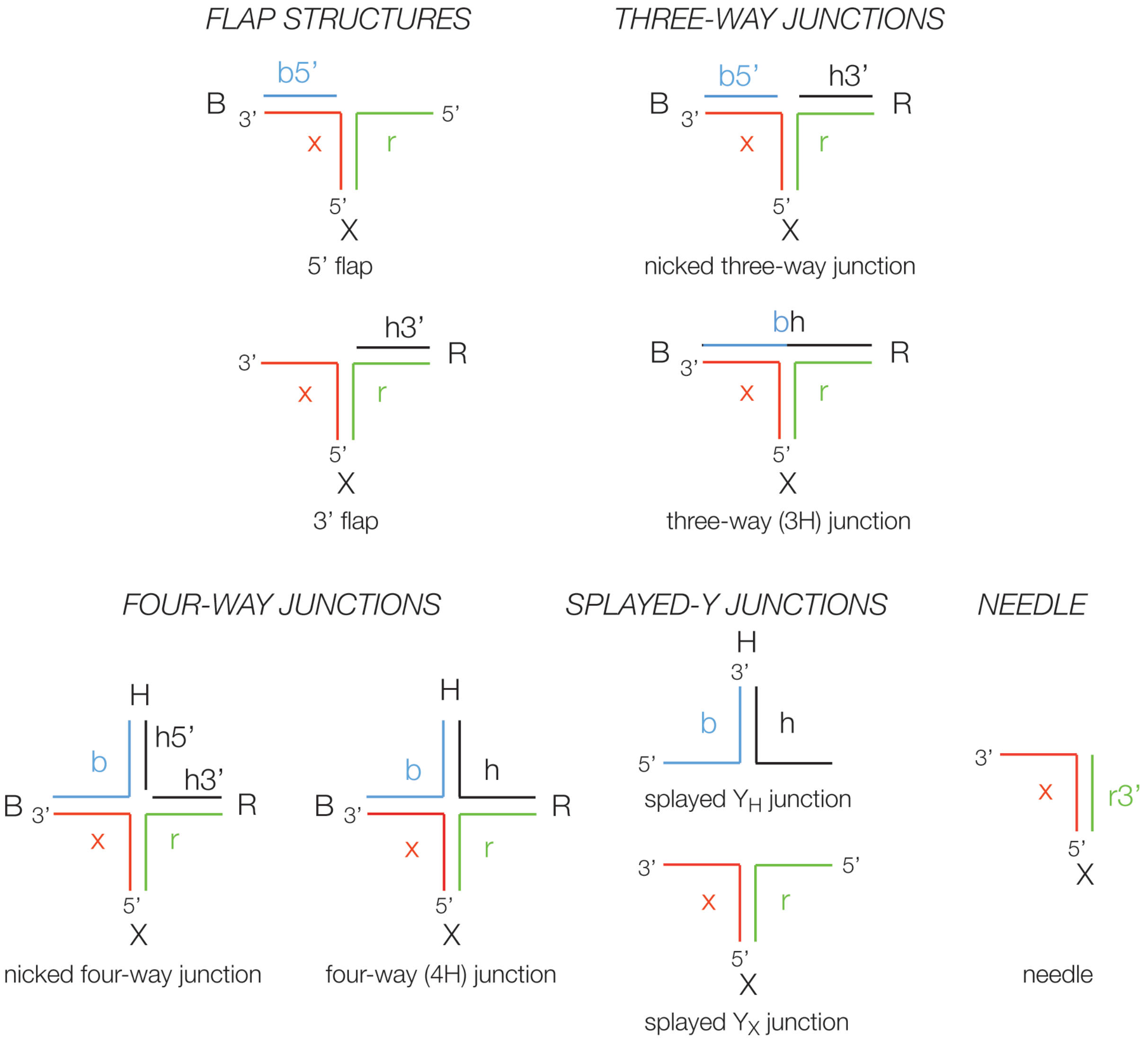


**Appendix Figure S4.** Relationship between the DNA junctions used as substrates for ANKLE1 in this work. All the junctions may be derived from the parental four-way junction 3, comprising four strands called b, h, r and x. These generate the four helical arms B, H, R and X named from the component 5' strand. The other junctions are generated by hybridising sub-sets of these strands, half-strands (e.g. *b5'* in the 5' flap structure) or composite strands (e.g. *bh* in the three-way junction). In the majority of the experiments we have used a radioactively-[5'-$^{32}$P] labelled x strand, shown red here. The sequences of the strands are tabulated in Table S1.

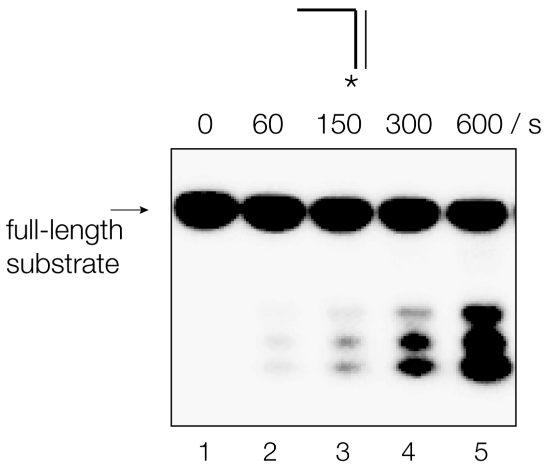


**Appendix Figure S5.** Cleavage of needle DNA structure by human ANKLE1. This simple structure is cleaved by hANKLE1, but at a rate that is slower than that of the splayed Y junction.

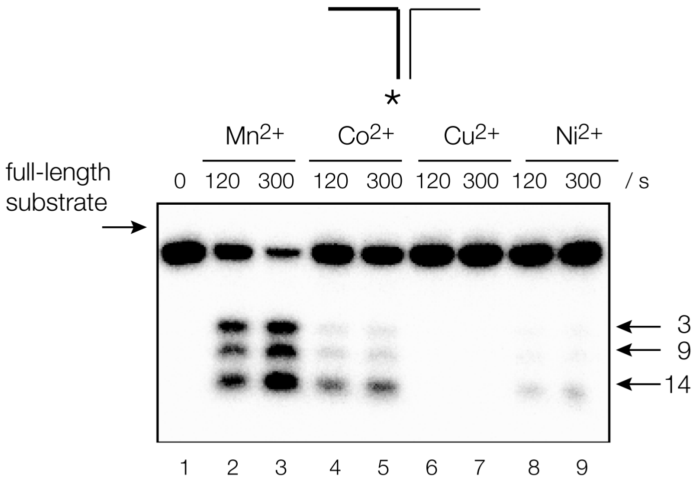


**Appendix Figure S6.** Cleavage of splayed $Y_X$ DNA junction by human ANKLE1 as a function of the divalent metal ion present. Radioactively [5'-$^{32}$P]-labelled DNA was incubated with hANKLE1 in the presence of 20 mM cacodylate (pH 6.5), 50 mM KCl, 0.1mg/ml BSA and 2 mM indicated divalent metal ion chloride for 120 and 300 seconds. Very little activity was observed in the presence of $Mg^{2+}$ or $Ca^{2+}$ ions.

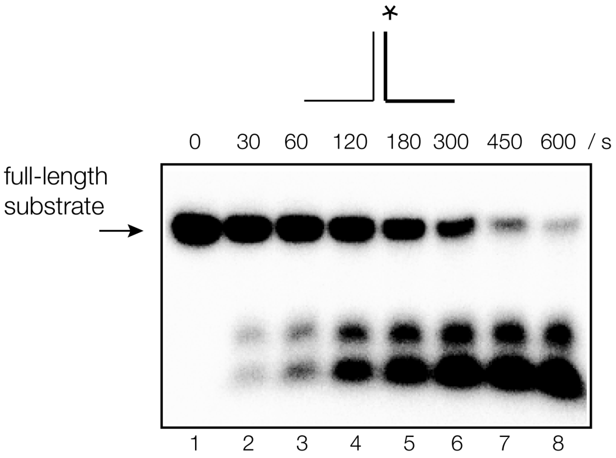


**Appendix Figure S7.** Cleavage of a different splayed-Y junction by human ANKLE1. The splayed $Y_H$ junction is derived from the H arm of the four-way junction 3 (see Figure S4). Radioactively [5'-$^{32}$P]-r-strand labelled DNA was incubated with hANKLE1 under single-turnover conditions. Despite having a completely different sequence from the splayed $Y_X$ DNA, it is nevertheless a good substrate for hANKLE1.

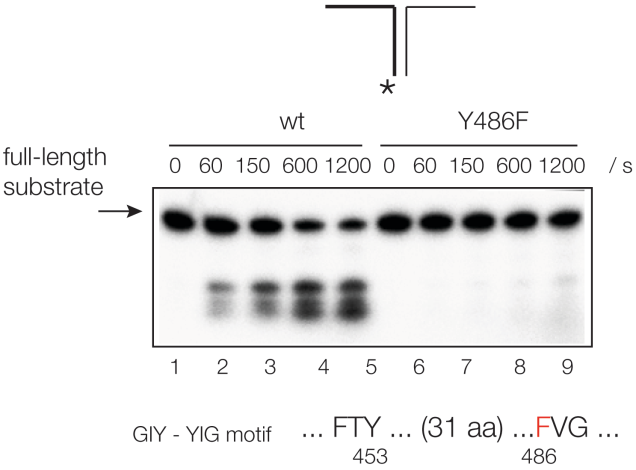


**Appendix Figure S8.** Mutation of tyrosine 486 in the GIY-YIG motif prevents cleavage of a splayed-$Y_X$ DNA junction by human ANKLE1. This experiment was performed using a GST-hANKLE1 (615 amino acid) fusion. Radioactively [5'-$^{32}$P]-labelled splayed $Y_X$ junction DNA was incubated with hANKLE1 under single-turnover conditions.

### SUPPLEMENTARY TABLES

b-strand :

CCTCGAGGGATCCGTCCTAGCAAGG GGCTGCTACCGGAAGCTTACAGATG

h-strand :

CATCTGTAAGCTTCCGGTAGCAGCC TGAGCGGTGGTTGAATTCACAGATG

r-strand :

CATCTGTGAATTCAACCACCGCTCA ACTCAACTGCAGTCTAGAACACATG

x-strand :

CATGTGTTCTAGACTGCAGTTGAGT CCTTGCTAGGACGGATCCCTCGAGG

bh-strand :

CCTCGAGGGATCCGTCCTAGCAAGG TGAGCGGTGGTTGAATTCACAGATG

b5'-strand :

CCTCGAGGGATCCGTCCTAGCAAGG

h5'-strand :

CATCTGTAAGCTTCCGGTAGCAGCC

h3'-strand :

TGAGCGGTGGTTGAATTCACAGATG

r3'-strand :

ACTCAACTGCAGTCTAGAACACATG

Complementary r-strand (used to generate double-stranded DNA) :

CATCTGTGAATTCAACCACCGCTCAGGCTGCTACCGGAAGCTTACAGATG

**Appendix Table S1.** The sequences of the oligonucleotides used in the construction of the hANKLE1 substrates studied in this work. All sequences are written 5' to 3'. A gap has been left at the point of strand exchange a junction is constructed from a combination of these oligonucleotides. The combinations of strands used to generate the different species are shown in Figure S4.

| plasmid | vector | description | UniProt accession number |
|---|---|---|---|
| JF1 | pKL | GST tagged hANKLE1669aa | A0A499FJM0 |
| JF2 | pKL | GST tagged hANKLE1615aa | Q8NAG6 |
| JF3 | pKL | GST tagged hANKLE1615aa with a point mutant, Y453F | Q8NAG6 |
| JF4 | pKL | GST tagged hANKLE1615aa with a point mutant, Y486F | Q8NAG6 |

**Appendix Table S2.** The plasmids used to express ANKLE1 in these studies.
